# Dual CRISPR-Interference Strategy for Targeting Synthetic Lethal Interactions between Non-Coding RNAs in Cancer Cells

**DOI:** 10.1101/2024.11.05.622038

**Authors:** Jeffrey L. Schloßhauer, Sama Shamloo, Katja Schamrin, Jochen Imig

## Abstract

Long non-coding RNAs (lncRNAs) represent a vast and functionally diverse class of RNA molecules, with over 100,000 predicted in the human genome. Although lncRNAs are less conserved across species compared to protein-coding genes, they play critical roles in gene regulation, chromatin interactions, and cancer progression. Their involvement in cancer make them promising therapeutic targets. CRISPR interference (CRISPRi), utilizing catalytically inactive Cas9 fused with a transcriptional repressor such as KRAB-MeCP2, offers a precise method for targeting nuclear lncRNAs and assessing their functions. This study introduces a dual CRISPRi system using orthogonal CRISPRi technologies from *Staphylococcus aureus* and *Streptococcus pyogenes* dCas9-KRAB, optimized for combinatorial targeting of lncRNAs in human melanoma cells. The protocol facilitates combinatorial gene knockdown or synthetic lethal screening of lncRNA pairs, providing a novel tool for cancer research. By exploring synthetic lethality between lncRNAs, this approach can help identify lncRNA interactions critical for cancer cell survival, offering new therapeutic strategies. The dual system’s functionality is demonstrated, highlighting its potential in identifying critical cancer-specific lncRNA interactions.

**SUMMARY:** This study presents a dual CRISPRi system targeting long non-coding RNAs in melanoma cells. It enables combinatorial gene knockdown and synthetic lethal screening, identifying cancer-specific lncRNA interactions for potential therapeutic strategies.

## INTRODUCTION

Although less than 3% of the human genome encodes proteins, approximately 80% of the genome is transcribed^1,2^. Among the non-coding transcriptional units, tens of thousands are classified as long non-coding RNAs (lncRNAs) exceeding 200 nucleotides, and the total number of human lncRNAs is estimated to exceed 100,000^3,4^. In contrast to coding genes, lncRNAs are less conserved across species. Although humans share 99% of their genome with primates like chimpanzees, lncRNAs are hypothesized to have a much greater influence on phenotypic evolution^5,6^. These findings indicate important cellular functions of lncRNAs. Although the regulation of lncRNAs and their interactions with RNA-binding proteins and other RNAs remain incompletely understood, and many lncRNAs are yet to be fully annotated, it is clear that lncRNAs exhibit cell- and tissue-specific expression patterns in health and disease, such as cancer^7–10^. They are implicated in diverse functions, including gene transcription regulation, involvement in chromatin interactions^11^, RNA processing^12^, RNA stabilization^13^, and regulation of translation^14^.

In cancer, diverse highly cell type-specific lncRNAs influence tumor development and metastasis by regulating gene expression, highlighting their potential as valuable therapeutic targets^15^. Beyond detection of lncRNAs as biomarkers in tumor samples^16^, targeting tumor-specific lncRNAs to disrupt their downstream functions holds significant potential in both clinical applications and basic research. RNA-based approaches to elucidate the roles of lncRNAs include antisense oligonucleotides (ASOs), short hairpin RNAs, and small interfering RNAs (siRNAs)^17,18^. While siRNA is commonly utilized for gene silencing screens, the siRNA-based knockdown is restricted to the cytoplasm^19^. However, lncRNA frequently operate within the nucleus.

Alternatively, Clustered Regularly Interspaced Short Palindromic Repeats interference (CRISPRi) can be used to inhibit lncRNAs in human cancers^20^. Moreover, genome-wide CRISPRi screens can be easily programmed and target a wide range of coding and non-coding genes to examine their functional impact^21,22^. In CRISPRi a catalytic deficient Cas9 (dCas9) is fused to a transcriptional repressor domain, such as the Krüppel-associated box (KRAB) domain^23^. Gene repression by dCas9-KRAB is guided by a guide RNA (gRNA) to the region of interest. CRISPRi controls genes at the DNA levels leading to higher efficiency and desired loss-off-function phenotypes in contrast to RNA interference, which is active at the post-transcriptional level^24^. In response to the limited efficacy of KRAB in target silencing, a fusion of KRAB and MeCP2 with dCas9 was introduced as a more effective repressing strategy^25^.

Although single gene silencing may affect cancer viability, synthetic dual or multiple lethal interactions can rescue cancer cells from cell death^26^. Synthetic lethality involves two or more genes, which can compensate the function of the other one. To overcome issues with single-gene knockdown screens, dual CRISPR strategies targeting lethal protein coding gene pairs offer a promising approach^27^.

Here, we present a protocol for the combinatorial use of orthogonal CRISPRi-based targeting of lncRNA or other non-coding RNA pairs using *Staphylococcus aureus* and *Streptococcus pyogenes* dCas9 fused to KRAB in human melanoma cells (Figure 1). The protocol can be utilized for combinatorial classical CRISPR knockdown or as CRISPRi-based screening of synthetic lethal pairs in cancer. In conclusion, we provide a fully functional dual CRISPRi system in melanoma cells as a cancer model.

**Figure 1:**
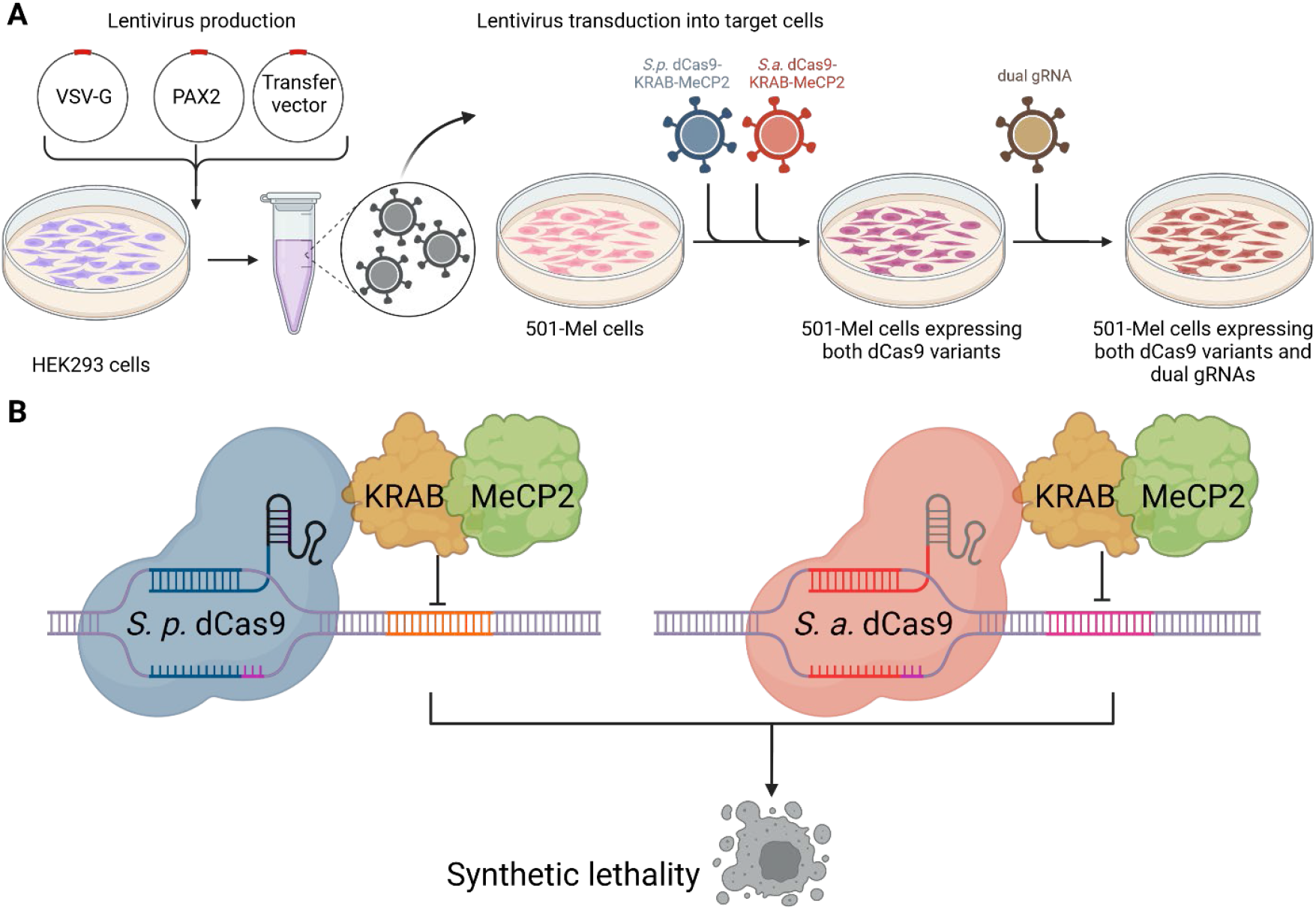
Scheme of the dual CRISPRi system to target synthetic non-coding RNA lethal interactions. **(A)** HEK293 cells were transfected with envelope plasmid VSV-G, packaging plasmid PAX2, and transfer vector to produce lentiviruses for subsequent transduction into 501-mel cells. Lentiviruses containing the genetic information for *Sp*dCas9-KRAB-MeCP2 (blue) and *Sa*dCas9-KRAB-MeCP2 (red) were integrated into 501-mel cells simultaneously. Following antibiotic selection, a second lentiviral transduction was performed to integrate the desired dual gRNAs (olive green). **(B)** 501-mel cells expressing the dCas9 variants interact with their corresponding gRNAs to silence target gene expression, resulting in cancer cell death. Created with BioRender.com.

## PROTOCOL

The human melanoma cell line 501-mel (RRID: CVCL_4633) was kindly provided by the Aifantis Lab (New York University). The Lenti-X 293T HEK cell line was purchased from Takara Bio. These cell lines were cultured in Gibco Dulbecco’s modified Eagle’s medium (DMEM) high glucose supplemented with 10% fetal bovine serum (FBS) (Sigma Aldrich) at 37 °C in a 5% CO_2_ atmosphere under sterile conditions. Cells were passaged until they reached 80-90% confluence.

### 1. LncRNA selection and gRNA design

The gRNAs were designed using Benchling R & D Cloud platform (retrieved from https://benchling.com), following site-specific recommendations to prioritize the highest on-target activity while minimizing off-target matches in both cases. Positioning of the gRNAs was done into a window of -150-+ 50 bp around the transcriptional start site (TSS) for most optimal performance. The *S. pyogenes* gRNAs were extracted from Petroulia *et al*.^28^. The designed gRNAs are listed in Supplementary Table 1. The lncRNAs RP11-120D5.1 (referred to as RP11 for convenience) and XLOC_030781 (referred to as XLOC for convenience) were targeted with one gRNA corresponding to *S. pyogenes* (gRNA-sp, the top-performing guide from the library screen) and three gRNA corresponding to *S. aureus* (gRNA-sa). A non-targeting control against the Rosa26 was combined with each target gRNA to create the dual gRNA vector, leading to knockdown one target lncRNA. Simultaneous knockdown of both, RP11 and XLOC, was achieved by exchanging the control gRNA Rosa26 in the dual gRNA vector by the equivalent target gRNA.

### 2. gRNA cloning

An overlap extension PCR was carried out using two DNA fragments containing the gRNA sequence. The U6 forward primer and H1 reverse primer (Supplementary Table 2) were employed for amplification in analogy to Najm *et al*.^27^.

1. A 50 µL PCR reaction was set up, consisting of 25 µL of 2X Phusion High-Fidelity PCR Master Mix with HF Buffer and 1 µL of each DNA fragment containing either the corresponding *S. pyogenes* or *S. aureus* gRNA. The reaction was run under the following cycling conditions: initial denaturation at 98 °C for 30 seconds, followed by 15 cycles of denaturation at 98 °C for 5 seconds, annealing at 55 °C for 10 seconds, and elongation at 72 °C for 15 seconds.
2. After completing the initial 15 cycles without primers, 2.5 µL of U6 forward primer and 2.5 µL of H1 reverse primer were added, and the reaction was subjected to an additional 20 cycles under the same conditions with a final extension for 5 min at 72 °C.
3. The PCR products were purified by the QIAquick PCR Purification Kit according to the manufacturer’s instructions.
4. The modified plasmid pPapi-dual-gRNA-Zeo-GFP (modified from Addgene #96921, Supplementary Figure 1) was digested by the restriction enzyme Esp3I (BsmB1) according to the manufacturer’s instructions.
5. The PCR products were cloned into the digested pPapi-dual-gRNA-Zeo-GFP plasmid according to the NEBuilder HiFi DNA Assembly Reaction Protocol.
6. Chemical competent One Shot Stbl *E. coli* cells were transformed by mixing 50 µl bacteria with 4 µl assembly product according to the manufacturer’s instructions.

### 3. Lentivirus Production

CAUTION: You are working with active lentiviruses at all these steps; follow the appropriate safety guidelines.

1. 4 × 10^6^ Lenti-X 293T HEK cells per 10 cm plate were plated to achieve ∼80% confluency the next day.
2. On the day of transfection, 500 μL Opti-MEM was mixed with 11.25 µg transfer vector *S. pyogenes* dCas9-KRAB-MeCP2 (SpdCas9-KRAB-MeCP2) (11.25 μg), 5.5 µg envelope plasmid VSV-G (Addgene #8454), and 16.5 µg of pPAX2 vector (Addgene #12260). This mixture was incubated for 5 min at room temperature. NOTE: Each lentivirus batch was produced using a distinct transfer vector. For the second dCas9 variant, the transfer vector utilized was *S. aureus* dCas9-KRAB-MeCP2 (SadCas9-KRAB-MeCP2). For the dual gRNA experiments, the transfer vectors were designed to target either the *RP11* locus (with gRNAs RP11-sp and RP11-sa3) or the *XLOC* locus (with gRNAs XLOC-sp and XLOC-sa3), depending on the species. These were combined with a control gRNA targeting the *Rosa26* locus. To target both loci simultaneously, transfer vectors were constructed with combinations of RP11-sp and XLOC-sa3, or XLOC-sp and RP11-sa3.
3. In a separate tube, 500 μL of Opti-MEM was mixed with 36 μL PEI reagent (stock: 1 mg/mL) to obtain a 1.5:1 ratio of PEI to total DNA. This mixture was incubated for 5 min at room temperature.
4. Both mixtures were combined and incubated for 15 min at room temperature.
5. The mixture was added dropwise to the cell medium (6 mL), and the cell plates were incubated overnight at 37 °C and 5% CO_2_. NOTE: Cells should not be allowed to be detached, this will decrease lentiviral yield.
6. After 12-15 hours, the medium was replaced with 5 mL of fresh DMEM medium.
7. Lentivirus-containing supernatant was harvested 48 hours post-transfection and replaced with fresh medium. Second and third harvests occurred 72 and 96 hours post-transfection, respectively. The supernatants were stored at 4 °C until final harvesting.
8. All supernatants were combined and centrifuged for 5 min at 500 x g and 4 °C to pellet detached cells and debris.
9. The supernatant was filtered through 0.2 μM sterile cell culture grade syringe filters.
10. The filtered supernatant was concentrated at 1000 x g and 4 °C using a Amicon Ultra-15 Centrifugal Filter Unit with a 100 kDa cut off to around 500 μL.
11. Virus aliquots of 50-100 µL were stored at -80 °C until further use. NOTE: For cell types that are challenging to transduce with lentivirus, such as suspension cells, the transfection process can be proportionally scaled up to 1–2 15 cm plates. Ultracentrifugation may be used to further concentrate the lentiviral supernatant. Target cells can then undergo spinoculation by centrifuging at 1500 rpm for 2–3 hours at room temperature, using a plate insert.

### 4. Lentivirus transduction and selection of stable transfected cells

CAUTION: You are working with active lentiviruses at all these steps; follow the appropriate safety guidelines under biosafety level 2 laboratory environment.

1. 501-mel cells were plated in a 6-well plate at 2 × 10^5^ cells/well with a final culture volume of 2 mL one day before transduction.
2. Culture medium was exchanged by 2 mL prewarmed DMEM medium supplemented with polybrene at a final concentration of 6 μg/mL.
3. The prepared lentiviruses were added to the cells dropwise. The plate was shaken slowly and incubated at 37 °C in a 5% CO_2_ atmosphere. NOTE: 50 µL of lentiviruses containing the information for SpdCas9-KRAB-MeCP2 and SadCas9-KRAB-MeCP2 were added simultaneously to the cells.
4. Sixteen hours post-transduction, the medium was replaced with fresh medium containing the selection antibiotics blasticidin and puromycin at a final concentration of 10 µg/mL and 2 µg/mL respectively.
5. Cells were selected until all control cells, which were not transduced, had died, with medium exchanged every second day. NOTE: Alternative antibiotic selection strategies are possible, depending on the resistance cassette in the transfer vector.

### 5. Protein quantification and western blot analysis

1. 10^6^ stable 501-mel transfected cells were centrifuged for 5 min, 500 x g at 4 °C.
2. The supernatant was discarded and the cell pellet resuspended with 1 mL ice-cold PBS.
3. Centrifugation was repeated and the cell pellet was resuspended in 50 µL Radio-immunoprecipitation assay (RIPA) buffer, containing 25 mM Tris-base, 150 mM NaCl, 1% NP-40, 0.5% sodium deoxycholate, 0.1% sodium dodecyl sulfate (SDS), 1x cOmplete Protease Inhibitor Cocktail.
4. After incubating on ice for 15 min, cell debris and insoluble material were removed by centrifugation at 13,000 × g for 10 min at 4 °C. The supernatant was either stored at -20 °C for future use or directly used for protein quantification using the Pierce BCA Protein Assay Kit according to the manufacturer’s instructions.
5. A total of 10 µg of protein was separated by size using SDS-PAGE in the Mini-PROTEAN Tetra Handcast System.
6. The resolved proteins were transferred to an Immobilon-P PVDF membrane using the Mini Trans-Blot Cell system.
7. The membrane was incubated for one hour at RT in blocking buffer (5% non-fat milk powder in TBST) with gentle shaking at 80 rpm.
8. The primary antibodies anti-Cas9 (*S. aureus*) (6H4) mouse monoclonal antibody and anti-Cas9 (*S. pyogenes*) (7A9-3A3) monoclonal antibody were added separately to the membrane at a 1:1,000 dilution in blocking buffer. The anti-GAPDH mouse monoclonal antibody was utilized as internal control and incubated with the membrane at a 1:1,000 dilution in blocking buffer.
9. After incubating over night at 4 °C with gentle shaking at 80 rpm, the membrane was washed three times with TBST buffer for 10 min at RT.
10. A secondary anti-mouse HRP antibody was incubated at a 1:1,000 dilution with the membrane for one hour at RT and 80 rpm.
11. The membrane was washed three times with TBST buffer for 10 min at RT.
12. Signal detection was performed using the Clarity Western ECL Substrate according to the manufacturer’s instructions and the chemo luminescence signal was detected using the Bio-Rad ChemiDoc MP Imaging System.

### 6. Dual LncRNA repression viability assay

CAUTION: You are working with active lentiviruses at all these steps; follow the appropriate safety guidelines under biosafety level 2 laboratory environment.

1. 501-mel cells containing stable transfected SadCas9-KRAB and SpdCas9-KRAB were infected with lentiviruses containing the dual gRNA vectors 7 days following transduction with dCas9 variants, using the same conditions as previously outlined. NOTE: 100 µl lentiviruses containing the dual gRNA vector were added to 2 × 10^5^ cells/well in 6-well plate.
2. Sixteen hours post-transduction, the medium was replaced with fresh medium containing 500 µg/mL Zeocin to enrich positively transduced cells. Selection pressure was maintained by adding 5 µg/mL blasticidin and 1 µg/mL puromycin.
3. Luminescence detection was performed one and five days post-transduction using the CellTiter-Glo Luminescent Cell Viability Assay according to the manufacturer’s instructions.

## REPRESENTATIVE RESULTS

The expression cassettes for transfected *Sa*dCas9-KRAB-MeCP2 and *Sp*dCas9-KRAB-MeCP2 were integrated into 501-mel cells simultaneously using lentiviral transduction. Following the enrichment of positively transfected cells, samples were collected for western blot analysis to confirm the presence of the dCas9-KRAB-MeCP2 variants (Figure 2A).

**Figure 2:**
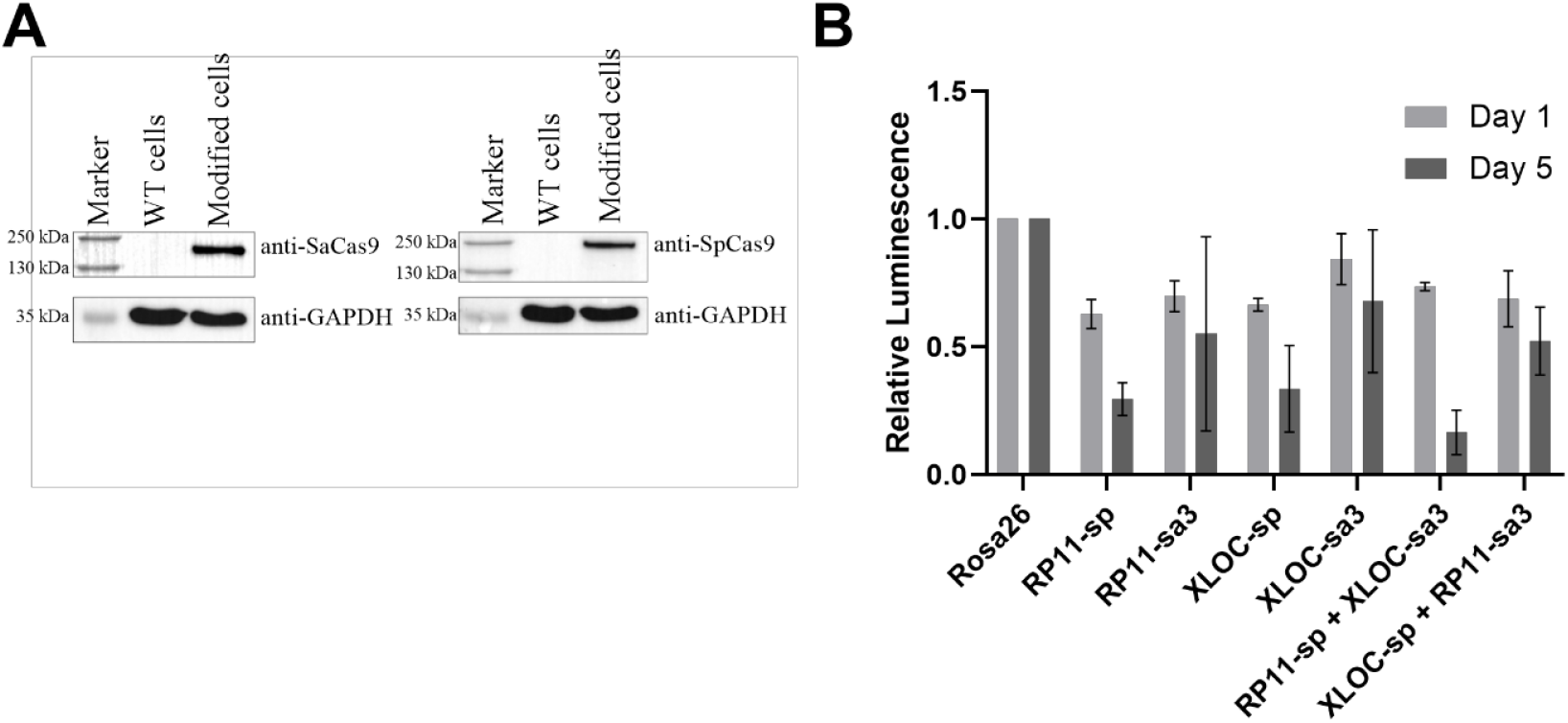
Functionality of the dual lncRNA repression system. **(A)** Western blot analysis was performed to detect the dCas9 variants in modified 501-mel cells. 501-mel wild type (WT) cells without transduction were used as control. **(B)** The dual lncRNA repression viability assay was performed after integrating the dual gRNAs into the modified 501-mel. Cells were analyzed after one and five days. Data are shown as mean ± SD.

Once 501-mel cells stably expressing the functional repressors were generated, a second lentiviral transduction was performed using a dual gRNA vector. RP11 and XLOC were previously shown to be upregulated in short-term cultures of metastatic melanoma, and individual CRISPRi targeting demonstrated their impact on proliferation and cell survival^28^. Thus, we selected this lncRNA pair for evaluation as a proof-of-concept synthetic lethal non-coding RNA combination, anticipating a cumulative, easily observable growth phenotype in the modified cells.

Different gRNAs targeting RP11 and XLOC were introduced to reduce the target mRNA levels. The repression of mRNA was investigated using target-specific gRNAs for either *Sa*dCas9-KRAB-MeCP2 or *Sp*dCas9-KRAB-MeCP2, tested individually and in combination, alongside a control gRNA targeting Rosa26.

As both XLOC and RP11 RNA levels were successfully reduced in the 501-mel cell line with the selected gRNAs, a dual lncRNA repression viability assay was performed using the most effective gRNA pairs (Figure 2B). After five days, a significant reduction in cell viability was observed with the combination of RP11-sp and XLOC-sa3, whereas only minor effects were seen when the target gRNAs were used individually.

## DISCUSSION

In this study, we implemented a dual lncRNA targeting strategy in melanoma cells using CRISPRi based on dCas9-KRAB-MeCP2 from two orthogonal species. Utilizing this system, we successfully repressed a synthetic lethal pair of lncRNAs, RP11 and XLOC, leading to cell death unlike when they were repressed individually.

The developed protocol is adaptable for targeting or screening other synthetic non-coding RNA lethal pairs by modifying the dual gRNA vector. Once a cell line of interest is stably expressing both dCas9 variants, a variety of approaches become feasible. While RNAi, ASOs, and shRNAs can be employed for gene silencing, CRISPRi offers a more straightforward screening methodology, and its combination with small chemical compounds or inhibitors is also conceivable. By substituting the dual gRNA vector with a dual gRNA library, this protocol allows for the systematic screening of synthetic lethal pairs in cancer cells, thereby enhancing the precision of cancer cell targeting.

Although we achieved integration of the dCas9-KRAB-MeCP2 variants through lentiviral transduction, alternative stable transfection methods can also be used to incorporate the desired enzymes into 501-mel cells. This is particularly relevant, as in our experience, the silencing of large constructs such as dCas9-KRAB-MeCP2 can occur over time. Therefore, consistent dCas9-KRAB-MeCP2 expression should be regularly monitored at recurring time points. Additionally, transducing two large fusion proteins along with selection markers and dual gRNA expression cassettes may induce cellular stress or may reach lentiviral packaging capacities.

Alternative strategies for integrating the CRISPRi system include the use of transposase vectors or CRISPR-directed integration to achieve stable insertion into “safe harbor” genomic sites, which are known for maintaining stable expression over time^29^. However, many cell types, including melanoma cells, are challenging to transfect^30^, making lentiviral transduction the preferred method. Electroporation of transposase- or Cas-based enzymes may offer an alternative means to integrate the dCas9-KRAB-MeCP2 constructs into melanoma cells^31^. Implementing an inducible knockout system would be advantageous for activating dCas9-KRAB-MeCP2 expression as needed, rather than constitutively, to minimize cellular stress.

Finally, the developed approach is not limited to melanoma cells but can be extended to any cell type where synthetic lethality is being investigated. This combinatorial versatility makes the protocol a valuable tool for exploring synthetic lethal interactions of lncRNAs across various cancer types and potentially other diseases.

## ACKNOWLEDGMENTS

We acknowledge Stefan Raunser (Max Planck Institute of Molecular Physiology, Dortmund) for access to Biosafety Level 2 lab space. Special thanks to Eric Wang for his intellectual contributions and to all past and present members of the Imig lab for their valuable discussions. Jochen Imig is currently CGCIII funded by Pfizer Inc. at CGC III.

