## Supplementary Information for "Dual CRISPR-Interference Strategy for Targeting Synthetic Lethal Interactions between Non-Coding RNAs in Cancer Cells"

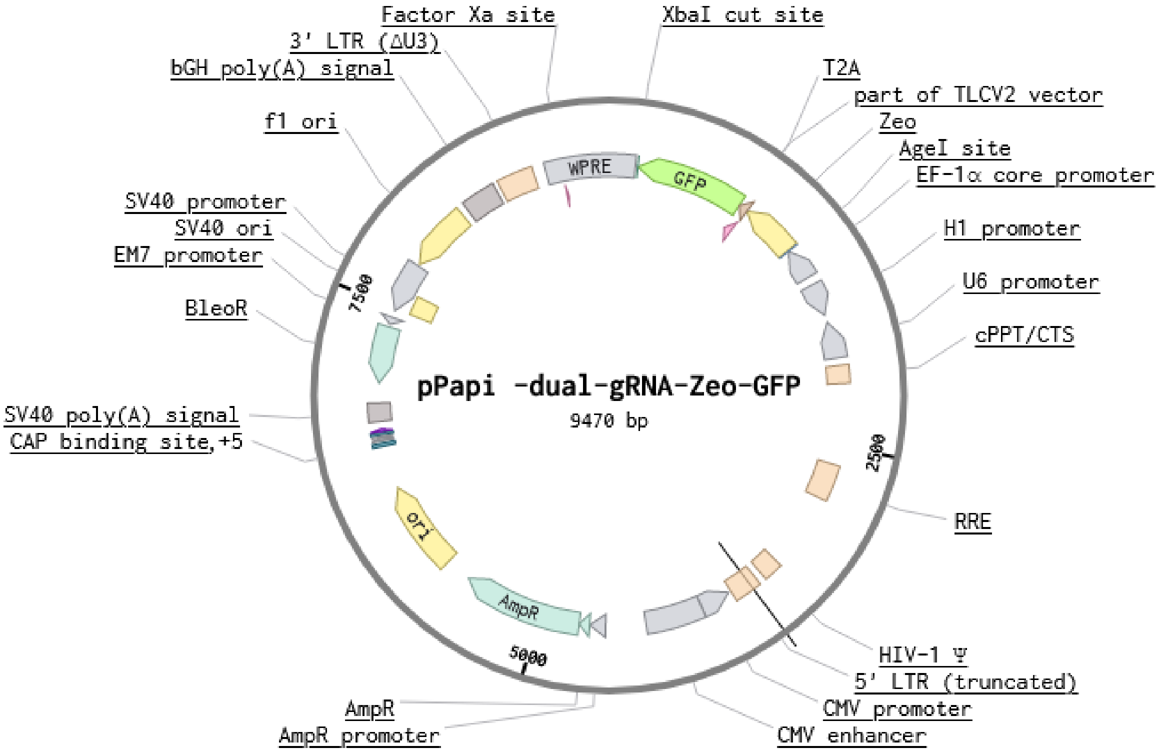


**Supplementary Figure 1: Plasmid map of pPapi-dual-gRNA-Zeo-GFP.** The plasmid was modified from Addgene #96921. The map was created with Benchling (https://benchling.com/).

**Supplementary Table 1: Utilized guide RNA sequences**

| **Name** | **Guide RNA sequence** | **dCas9** | **Target Gene** |
|---|---|---|---|
| RP11-s.p. | GGCGGTGACCGCAGCCACAG | S. pyogenes | RP11-120D5.1 |
| RP11-s.p.1 | TCTGCCGCATAAAGCATACA | S. aureus | RP11-120D5.1 |
| RP11-s.p.2 | GGGAAGCCTCTGGAGCAGAC | S. aureus | RP11-120D5.1 |
| RP11-s.a.3 | GAAACGTTAGTAGGTGCCCA | S. aureus | RP11-120D5.1 |
| XLOC-s.p. | GAGAAAGGAAGGAGGAAGGG | S. pyogenes | XLOC_030781 |
| XLOC-s.a.1 | GGAGTCTAACATACAAATAT | S. aureus | XLOC_030781 |
| XLOC-s.a.2 | ATTATTTAAAAAAAATTAAG | S. aureus | XLOC_030781 |
| XLOC-s.a.3 | GGAATTAGAACATGAACATT | S. aureus | XLOC_030781 |
| Rosa26 | AACGGCTCCACCACGCTCGG | S. pyogenes, S. aureus | Rosa26 |

**Supplementary Table 2: Utilized DNA sequences**

| Name | DNA sequence |
|---|---|
| U6 forward primer | GTGGAAAGGACGAAAcaccg |
| H1 reverse primer | GTATGAGACCACTCTTtcccg |
| DNA fragment S. pyogenes Rosa26 | GTGGAAAGGACGAAACACCGAACGGCTCCACCACGCT CGGGTTTGAGAGCTAGAAATAGCAAGTTCAAATAAGG CTAGTCCGTTATCAACTTGAAAAAGTGGCACCGAGTCG GTGCTTTTTTGAACCGACGGATGATCTCGTGCAC |
| DNA fragment S. aureus Rosa26 | GTATGAGACCACTCTTTCCCGAACGGCTCCACCACGCT CGGGTTTAAGTACTCTGGAAACAGAATCTACTTAAACA AGGCAAAATGCCGTGTTTATCTCGTCAACTTGTTGGCG AGATTTTTTGAACCGGTGCACGAGATCATCCGT |
